# Organelle bottlenecks facilitate evolvability by traversing heteroplasmic fitness valleys

**DOI:** 10.1101/2021.01.28.428572

**Authors:** Arunas L. Radzvilavicius, Iain G. Johnston

## Abstract

Bioenergetic organelles – mitochondria and plastids – retain their own genomes, and these organelle DNA (oDNA) molecules are vital for eukaryotic life. Like all genomes, oDNA must be able to evolve to suit new environmental challenges. However, mixed oDNA populations can challenge cellular bioenergetics, providing a penalty to the appearance and adaptation of new mutations. Here we show that organelle ‘bottlenecks’, mechanisms increasing cell-to-cell oDNA variability during development, can overcome this mixture penalty and facilitate the adaptation of beneficial mutations. We show that oDNA heteroplasmy and bottlenecks naturally emerge in evolutionary simulations subjected to fluctuating environments, demonstrating that this evolvability is itself evolvable. Usually thought of as a mechanism to clear damaging mutations, organelle bottlenecks therefore also resolve the tension between intracellular selection for pure oDNA populations and the ‘bet-hedging’ need for evolvability and adaptation to new environments. This general theory suggests a reason for the maintenance of organelle heteroplasmy in cells, and may explain some of the observed diversity in organelle maintenance and inheritance across taxa.

## Introduction

Mitochondria and plastids are organelles that exist in populations in eukaryotic cells and perform vital bioenergetic, metabolic, and signalling tasks. The majority of cells in the majority of eukaryotes contain mitochondria; cells in photoautotrophs also contain plastids. Originally independent organisms, mitochondria and plastids retain their own genomes (mtDNA and ptDNA respectively) that encode vital bioenergetic apparatus. Mutations in these organelle DNA (oDNA) molecules can compromise cellular function with fatal consequences [1].

oDNA molecules exist in dynamic populations in eukaryotic cells [2, 3, 4]. Each cell may contain hundreds or thousands of oDNA molecules, and these molecules may not be genetically identical. A mixture of genetically different oDNA molecules in a cell is referred to as heteroplasmic. Recent evidence suggests that small-scale heteroplasmy, where each genome differs at a small number of loci from others, is common in mtDNA populations [5, 6]. Larger-scale mtDNA diversity (differences at many loci) exist in populations of organisms, giving rise to, for example, the geographical distribution of different mitochondrial haplogroups in human population [1, 7, 8, 9].

As with nuclear genomes, organelle genomes are subject to evolution in sequence and structure; for example, experimental work has highlighted how environmental pressures shape mtDNA populations through selection [9, 10, 7, 11], although the role of environmental selection in human populations remains debated [12, 13]. The ability to evolve in response to environmental change is often termed evolvability [14, 15] and is an essential aspect of life. A classical question is how biological systems resolve the perceived tradeoff between two desirable characteristics: evolvability (producing new phenotypes through mutations) and robustness (retaining the same phenotype in the face of mutations) [16, 15]. This tradeoff can be resolved, at the population level, if a ‘robust’ subset of a population occupies regions of genetic space that are robust and an ‘evolvable’ subset occupies other regions from which other phenotypes are mutationally accessible [17]. Such genetic spread can act to ‘hedge bets’ against future environmental changes, by increasing the number of potential new solutions to uncertain future challenges [18, 19].

To evolve and adapt, mutational changes must be acquired by cellular oDNA populations, and these changes must be inherited from parent organism to offspring organism [20]. There is a negligibly small probability that the same mutational change is acquired simultaneously by all oDNA molecules in a cellular population, so the acquisition of mutational change necessarily induces cellular heteroplasmy [21]. However, recent experimental work has shown that some instances of even limited heteroplasmy can compromise cellular and organismal function [22, 23, 24, 25]. Genetic differences in mtDNA molecules within a cell may lead to incompatibilities between their encoded protein products, reducing bioenergetic capacity [25]. This raises the question: how can potentially beneficial mutation changes be acquired by oDNA populations, if the corresponding heteroplasmy has a negative impact on fitness?

The inheritance of mtDNA in many organisms involves a genetic ‘bottleneck’, where the effective number of mtDNA molecules is reduced as the female germline develops [26, 1, 27, 28]. This bottleneck can arise in part, but not necessarily completely, from a physical reduction of mtDNA copy number, and from stochastic cell divisions, mtDNA turnover, and other processes [29, 30, 31,32, 33, 27]. For heteroplasmic mothers, this bottleneck increases cell-to-cell variability in heteroplasmy level in the germ cells of the next generation (and selection may act to change mean heteroplasmy level in parallel [34, 35, 36]. For example, a mother carrying 50% type A mtDNA and 50% type B mtDNA may produce egg cells with a range of 30-70% type A. This increased variability can then be exploited by purifying selection at the cellular level to remove cells with a high proportion of deleterious oDNA, helping to avoid Muller’s ratchet – the buildup of deleterious mutations over generations [37, 38, 39, 40, 41]. The bottleneck can allow rapid shifts of mtDNA makeup even between one generation and the next [42, 27].

In addition to helping clear deleterious mutations, the variability generated by the bottleneck has also been suggested to help adaptation [21]. Previous theoretical analyses have explored the role of cell-to-cell oDNA variability in fixing new mutants within model cells and populations [43, 44]. However, to our knowledge, the fitness penalty associated with heteroplasmic oDNA populations – and the consequent potential ‘barrier’ or fitness valley – has not yet been considered in this literature. Conversely, studies are beginning to explore the role of heteroplasmic costs on evolutionary and cellular behaviour [45, 46] but to our knowledge have yet to be linked to the bottleneck’s role in adaptation and purification. Whether heteroplasmic fitness penalties pose a theoretical barrier to oDNA adaptation thus remains to be determined.

Here, we asked whether the rapid shifting in oDNA type due to a genetic ‘bottleneck’ can help to overcome the fitness penalty associated with heteroplasmic oDNA populations, and thus enhance evolvability. Our corollary question was, if bottlenecks can help evolvability in this way, is robustness necessarily sacrificed? We proceed by introducing a simple model for cellular oDNA populations and their behaviour between generations. We then analyse this model with stochastic simulations to investigate the extent to which heteroplasmy variability facilitates the evolvability of oDNA populations, and whether bottlenecks and heteroplasmy can naturally emerge as evolutionarily positive ‘strategies’ where evolvability is essential.

## Model

We model cells in the female germline, or its equivalent, as containing a population of *n* oDNA molecules which can generally include a mixture of oDNA types. Each generation consists of *N* mother cells. A mother cell in one generation gives rise to *N_off_* daughter cells in the next generation. Our model does not assume that a generation involves a particular number of cell divisions (for example, around 25 in some mammalian germlines [47], or 1 in the case of unicellular organisms): the contribution of these divisions and other segregating processes are pooled in our segregation parameter. We rather consider cells at identical developmental points, one generation to the next. Our default parameterisation uses *N* = 20 and *N_off_* = 2 with *n* = 100; we verified that changes to this parameterisation had only limited influence on the structure of our results (Supplementary Figs. S1-S2). Each cell has a mutant proportion *h*, ranging between 0 and 1. Each daughter cell inherits an oDNA complement from its mother, with mutant proportion:

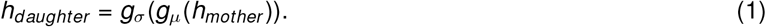

The functions *g_μ_* and *g_σ_* respectively describe changes in heteroplasmy due to mutation and segregation. We use 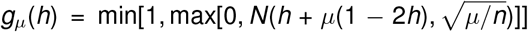, so that the mutant proportion in a cell is changed by a normal mutation kernel (*N*(*a, b*) denotes the normal distribution with mean a and standard deviation *b*), and constrained to lie between 0 and 1. The mean and variance of this kernel follow from the structure of the mutational processes that can act on the oDNA population (see Supplementary Information). We use 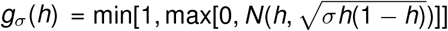, so that the variance induced by segregation is a quadratic function of initial mutant proportion (as observed in theory and experiment [27]), with final mutant proportion again constrained to lie between 0 and 1. The *σ* parameter in our model is then roughly comparable to the ‘normalised heteroplasmy variance’ *V’(h)* = *V*(*h*)/(*h*(1 - *h*)) commonly observed in genetic studies. For heteroplasmy distributions that approach normality, *σ* → *V’*(*h*); departures from normality make this equivalence weaker.

A cell has a fitness value with contributions from (i) the performance of its constituent oDNA molecules in the current environment, and (ii) an admixture penalty if the cell contains a mixture of different oDNA types. Each oDNA type provides a fitness contribution that is a function of environment, with different oDNA types performing better in different environments. The fitness function is sigmoidal, taking values between 0 and 1:

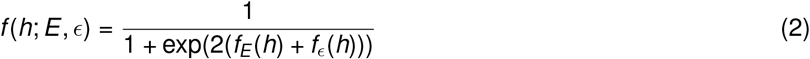

where *f_E_*(*h*) = -*h* in environment *A* and *h* in environment *B*, and *f_ϵ_*(*h*) = 0 for *h* = 0 and *h* =1, and -*ϵ* otherwise, following the observation that even limited heteroplasmy can challenge cell performance [25]. We also investigated a range of alternative fitness functions, including where the fitness was a smooth function of mutant proportion 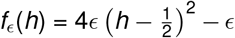, and where a ‘threshold effect’ keeps fitness constant until a given mutant proportion. This threshold effect models observations in mtDNA [48] where the severity of the phenotype is a nonlinear (even step-like) function of mutant proportion h.

Following assignment of fitness values, a new population of size *N* is selected from the *N* × *N_off_* daughter cells, with cells selected (with replacement) with probability proportional to their fitness (roulette wheel selection). Fitness and genetic content is tracked through *t_max_* generations (*t_max_* = 500 by default), and different regimes of environmental change, described for each experiment, are applied through generations.

The default picture above considers fitness to be governed by a single locus, so that mutant and wildtype oDNA molecules are the only types involved in the model. For generality we also consider a case where a third, dysfunctional, type of oDNA is included, and mutations can transform both mutant and wildtype oDNA molecules into this dysfunctional state. This models a picture where mutation leads to deleterious effects at other oDNA loci. In this case we define mutant proportion *h* = *m*/(*w* + *m*) (hence, the proportion of mutant oDNAs among the population of functional oDNAs), and introduce the dysfunctional proportion *h_D_* = *d*/(*w* + *m* + *d*), reporting the proportion of dysfunctional oDNAs in a cell. Cells are now described by {*h, h_D_*}, and the update rule now becomes 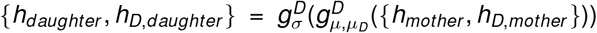, including a new mutation rate *μ_D_* = *α_μ_* that transforms functional to dysfunctional oDNA. Here, *α* reflects the relative propensity of mutations to switch between functional oDNA types versus inducing dysfunctionality. We derive the corresponding transition functions 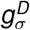 and 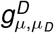 in the Supplementary Information, and redefine *f_E_*(*h, h_D_*) → *f_E_*(*h*) - *h_D_*, so that dysfunctional proportion has a purely detrimental effect on fitness. We show in the Supplementary Information that the main effect of this accounting for dysfunctional oDNA types is to limit the mutation rate *μ* for which viable populations of cells (those retaining some functional oDNA) can be maintained; below these mutational cutoffs, the interplay between mutation and bottlenecks remain comparable to the case without dysfunctional oDNA (Supplementary Figure S3).

## Results

### Heteroplasmy fitness costs challenge adaptation to new environments

We first asked to what extent a fitness penalty associated with heteroplasmy may challenge organelle adaptation to new environments. We initialised a model population of cells with homoplasmic organelle populations, all of the variant optimal for environment *A*. At time *t* = 0, we impose environment *B*, and track how organelle populations within the cell population subsequently evolve. We varied mutation rate *μ* and the magnitude of the fitness penalty *ϵ* experienced by cells with heteroplasmic populations.

We found that adaptation to the new environment was impossible in some circumstances, specifically for low mutation rates and high fitness penalties (Fig. 1A). Similar behaviour was observed for the range of alternative fitness functions we considered (Supplementary Fig. S4). In these regions, the selective penalty associated with the appearance of new mutations (and hence the appearance of heteroplasmy) prevents such mutations from ever propagating sufficiently to provide a fitness benefit from environmental matching. The penalty constitutes a ‘fitness valley’ that the system must overcome in order to fix new mutations. At higher mutation rates, the mutational load can increase rapidly enough – crossing the ‘valley’ – to allow this advantage.

**Figure 1:**
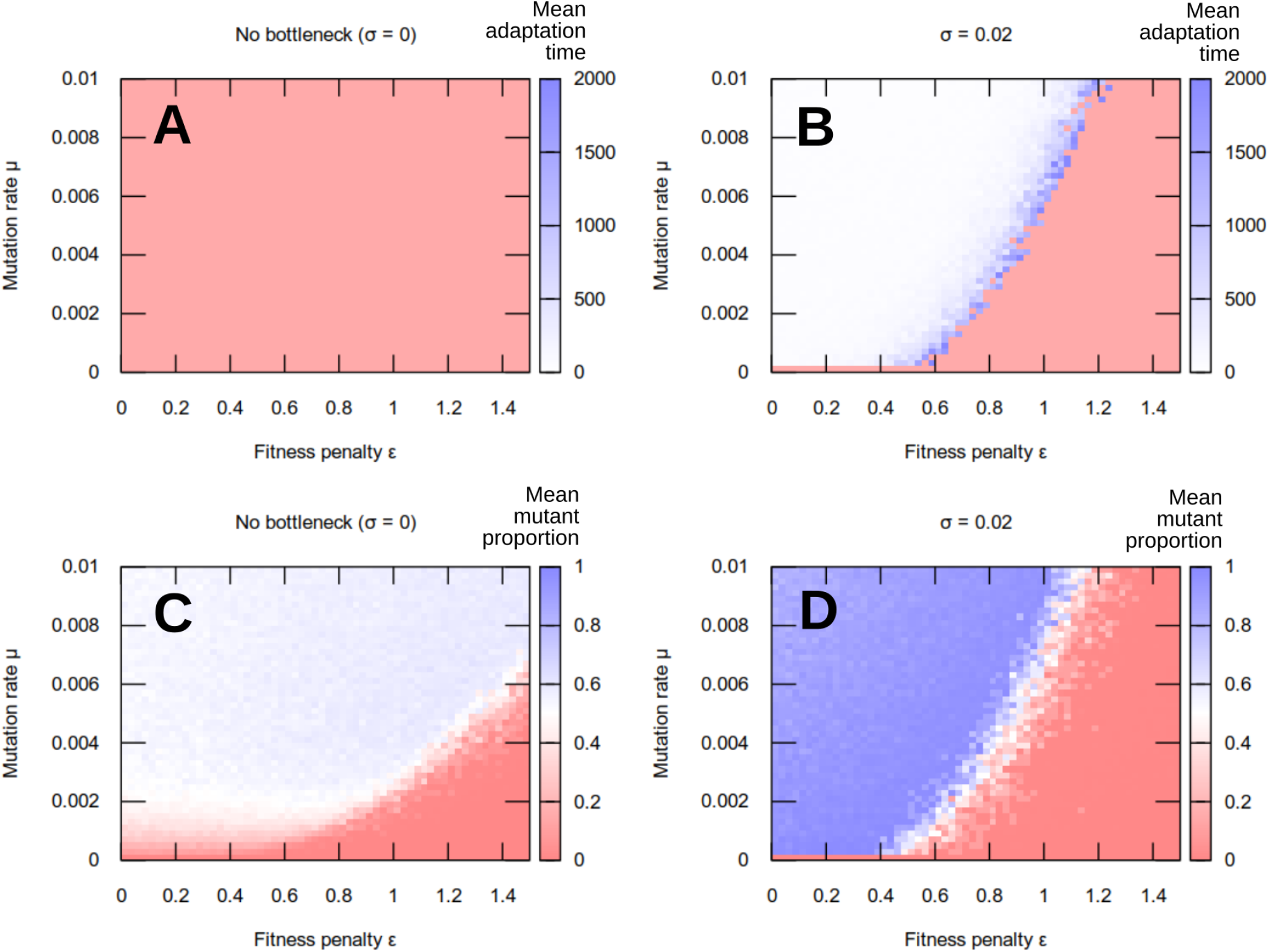
Adaptation to a new environment. (A, B) Adaptation times and (C, D) mean mutant frequency in a population after a change of environment, with heteroplasmy fitness penalty *ϵ* and mutation rate *μ*. (A, C) no between-generation segregation of mutations (*σ* = 0); (B, D) nonzero segregation (*σ* = 0.02). Without segregation, the ‘fitness valley’ prevents adaptation to the new environment, with the population either remaining adapted to the previous state (low mutation rate, high penalty) or reaching an intermediate state driven by mutation. Except in the case of zero mutation rate, segregation allows traversal of the fitness valley and facilitates adaptation.

### Segregation via the genetic bottleneck can overcome heteroplasmy fitness costs and allow adaptation to new environments

We next asked whether between-generation segregation of organelle mutations, via a genetic bottleneck, could help traverse this fitness valley and increase evolvability. We introduced this segregation via a new process in the model, where the mutant proportion of daughter cells differs from that of the mother cell by a normal random variate. This is not a perfect model of, for example, the mtDNA bottleneck, but captures the essential increase in variance in an intuitively simple way.

We asked how the magnitude of this segregation effect (the ‘tightness’ of the genetic bottleneck) influenced the system’s ability to adapt to new environments. The simulation setup was the same as in the previous section, with new variable *σ* describing segregation strength (Fig. 1A thus corresponds to *σ* = 0). We found that a nonzero segregation strength substantially enhances evolvability, allowing the propagation of beneficial mutations in regimes (particular those with low mutation rate and high fitness penalty) that posed the greatest challenge to adaptation without segregation (Fig. 1B). This behaviour was observed consistently across demographic and cellular parameterisations of our models, and when ‘off-target’ mutations inducing oDNA dysfunctionality was accounted for (Supplementary Figs. S1-S3).

### Robustness and evolvability with and without a bottleneck

Fig 1B shows that an organelle bottleneck increases evolvability of organelle populations. A parallel feature of populations is robustness – the ability to retain functionality in the face of mutational pressure. Traditionally, robustness and evolvability were viewed as opposites, but they can be reconciled at the population level [17].

In our model system, robustness can be pictured as related to the level of homoplasmy in the cellular population: the higher the proportion of the environmentally-optimal organelle type, the more mutation can be ‘absorbed’ without compromising the population. We found that the increased evolvability facilitated by organelle bottlenecks did not compromise system robustness: adaptation to new environments was achieved while long-term behaviour retained high levels of homoplasmy (Fig. 2). Because bottlenecks allowed adaptation at lower mutation rates, the bottleneck-adapted population was able to retain a more homoplasmic state than was possible when the population was forced to adapt through mutation alone. When dysfunctional mutations were likely, only low mutation rates were possible without losing population viability, but even small bottleneck sizes were sufficient to allow adaptation even at these low mutation rates (Supplementary Fig. S3).

**Figure 2:**
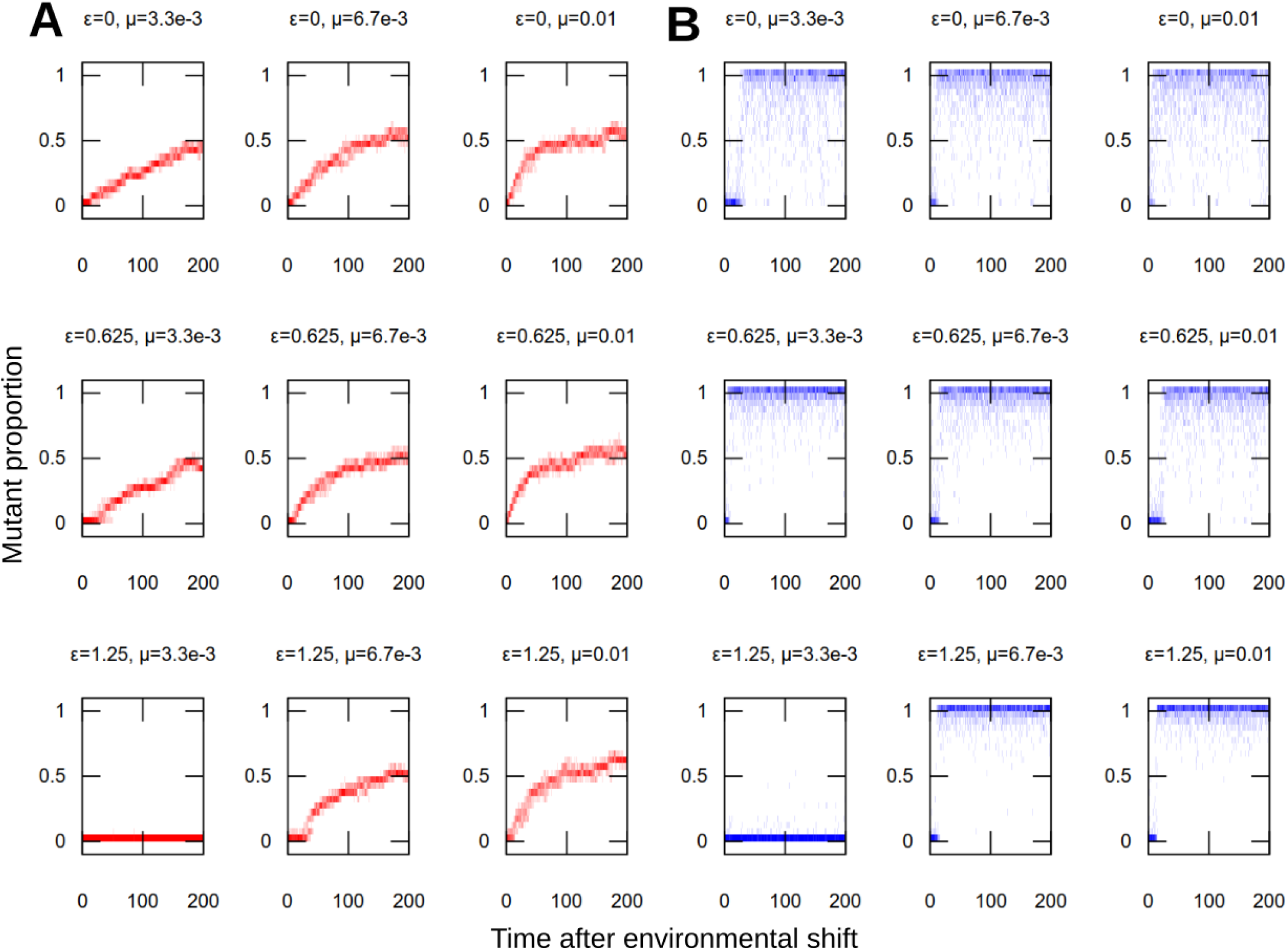
Population structure in adaptation to new environments. Population distribution of mutant proportion *h* with time in simulations adapting to a new environment (as in Fig. 1). (A) no segregation *σ* = 0; (B) nonzero segregation *σ* = 0.2. Distributions of *h* are presented for single example simulations with the given parameterisations. Lower mutation rates allow more homoplasmic populations (tighter *h* distributions), but, in the absence of segregation, challenge adaptation to the new environment (slow or no progress towards *h* = 1). Higher mutation rates facilitate adaptation but lead to more heterogeneous populations (wider *h* distributions). Segregation allows both rapid adaptation and high levels of homoplasmy.

### Interplay between mutation rate, fitness cost, and bottleneck strength

Following the observation that nonzero segregation enhances evolvability (without compromising robustness), we next asked how this enhancement depends on the magnitude of segregation strength (the ‘size’ of the genetic bottleneck). To this end we varied *σ*, the magnitude of segregation strength, and *ϵ*, the fitness penalty associated with heteroplasmy, for different values of *μ*, mutation rate (Fig. 3). Intuitively, for zero mutation rate, no adaptation was possible, because there was no generation of any genetic diversity at any time (Fig. 3A). However, for all nonzero mutation rates, almost any nonzero segregation immediately allowed a substantial enhancement of evolvability, with this enhancement then increasing more slowly as bottleneck size decrease further (Fig. 3B-D).

**Figure 3:**
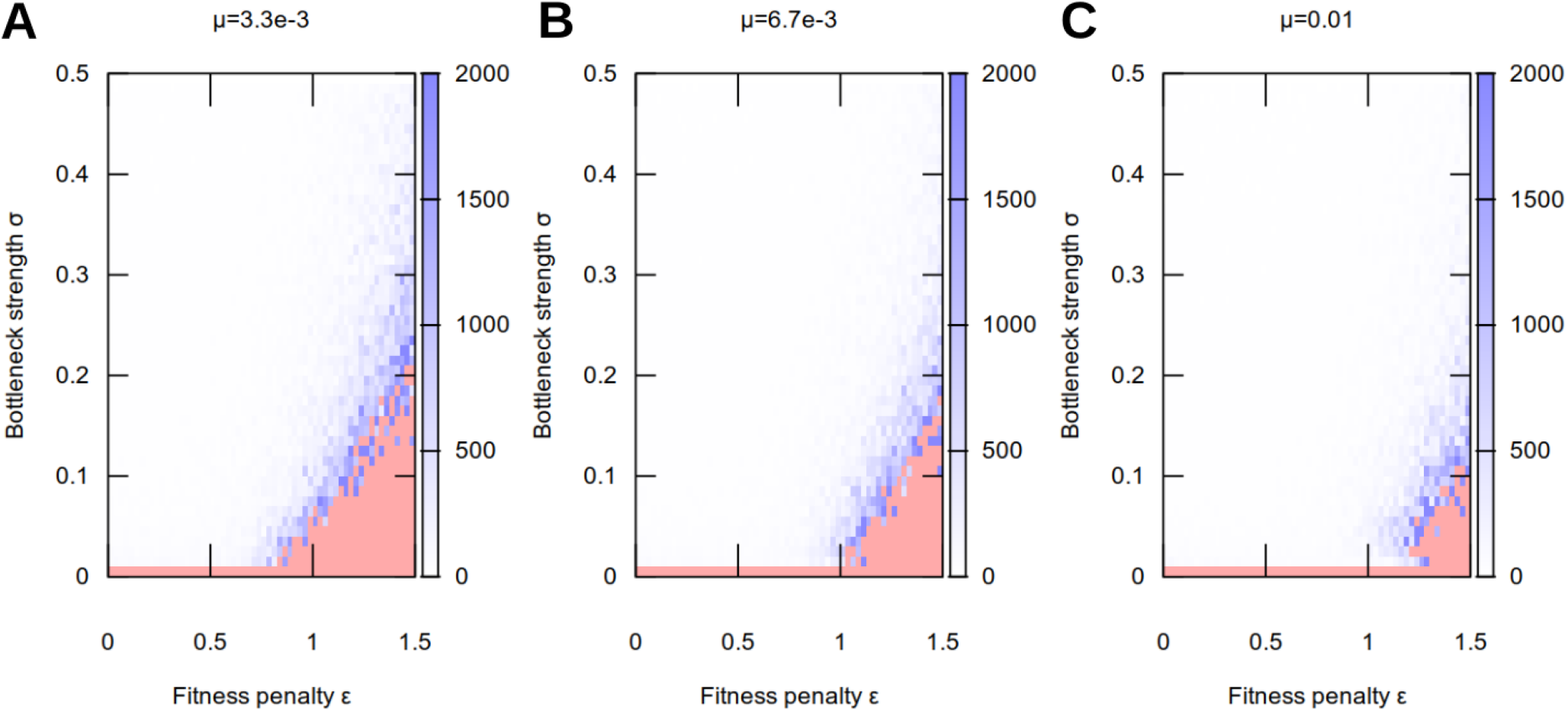
Adaptation to a new environment as a function of segregation strength. Plots show, as in Fig. 1, the mean time taken for a population to adapt following a change of environment. Increasing segregation strength *σ* facilitates adaptation at higher fitness penalties, with higher mutation rates *μ* amplifying this effect.

### Evolution and maintenance of mutation, bottlenecking, and costly heteroplasmy in changing environments

These results suggest that a combination of mutation and a ‘genetic bottleneck’ enhance evolvability in the sense of adaptation to new environments. To see if such a strategy could emerge in an evolving system, we next adapted our simulation so that mutation rate and ‘bottleneck size’ were themselves evolvable parameters within a population of cells. Specifically, we assign each cell in our model population a mutation rate *μ_i_* and a segregation strength *σ_i_*. These parameters are inherited from mother cell to daughter cells. This inheritance is imperfect, and a daughter’s mutation and segregation parameters can themselves mutate to be different from the mother. This ‘parameter mutation rate’ (different from the organelle mutation rate *μ_i_*) is labelled *δ*.

As the population of cells evolves, those cellular parameters that constitute evolutionarily successful strategies may thus come to dominate the population, while those parameters that constitute unsuccessful strategies will disappear. We simulated this system under different conditions. First, the new but then constant environment above (changing from environment A to environment B at time *t* = 0). Second, oscillating environments, changing from A to B with a period of *τ* generations. Lower values of *τ* correspond to faster environmental change; higher values of *τ* mean that environmental flips are infrequent. In each of these conditions, we observed the values of mutation rate and segregation strength that came to dominate the population.

We found that nonzero mutation rates and bottleneck strengths emerged in all conditions we consider (Fig. 4A-C). For static environments, their evolution was dependent on the magnitude of the fitness penalty. Lower fitness penalties allowed higher mutation rates and weaker bottlenecks; higher fitness penalties led to lower mutation rates and stronger bottlenecks.

**Figure 4:**
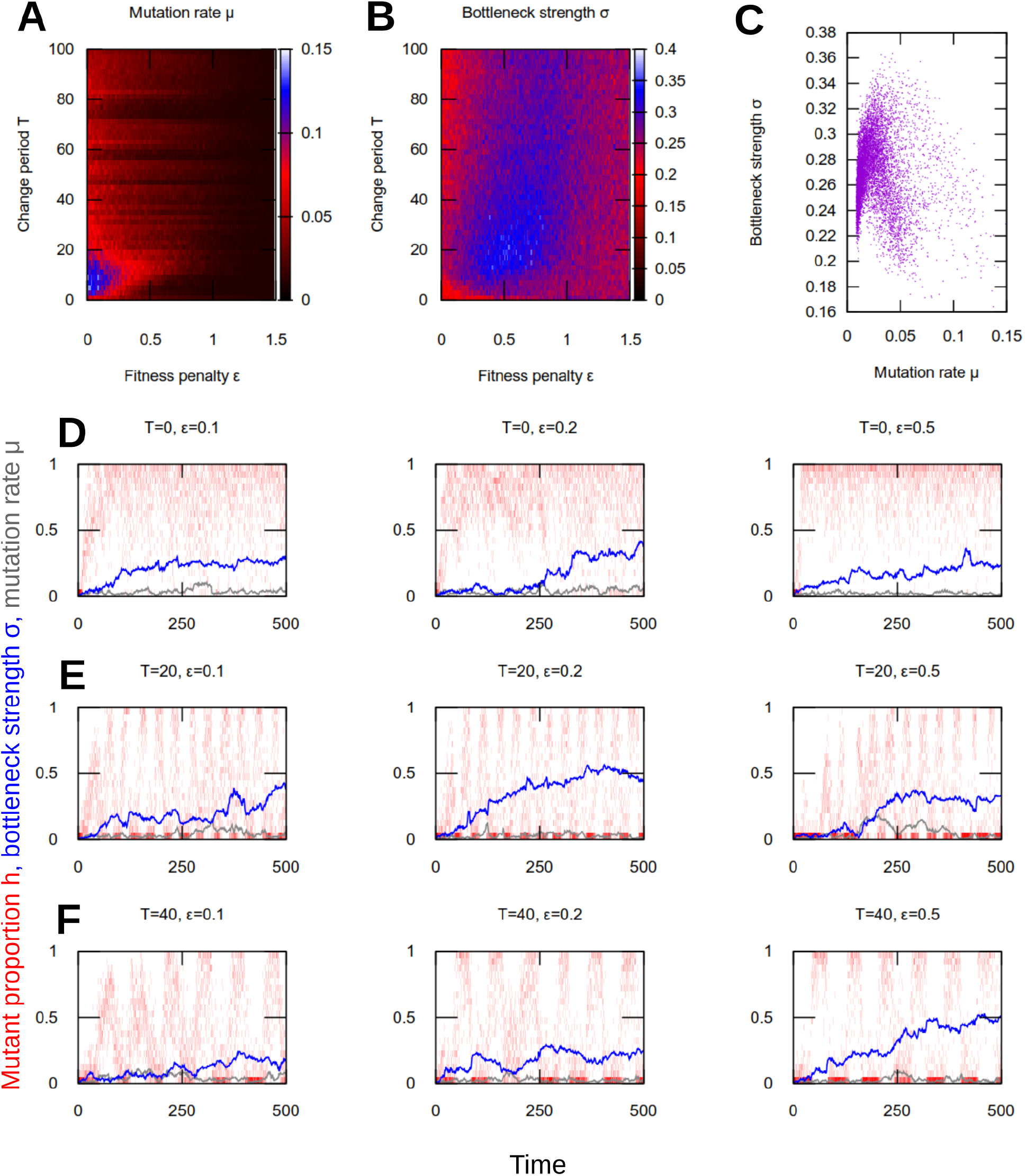
Evolution of mutation rate and segregation in dynamic environments. (A) mean mutation rate *μ*, (B) mean segregation strength *σ*, and (C) joint mean *μ* and *σ* after *t_max_* = 500 generations in evolutionary simulations (see text). Each point in (C) is from a particular parameterisation (i.e. a particular pixel in (A) and (B)). (D-F) Population histograms of mutant proportion *h* (red), plotted with traces of evolving mean mutation rate *μ* (grey) and segregation strength *σ* (blue), for different parameterisations and environmental change regimes: (D) no change; (E) rapid change; (F) slower change. Results are from single, representative simulations, many of which are averaged to give the results in (A-C). When environments change, mutation and segregation evolve together to facilitate adaptation (rapid shifting with environment) and robustness (high proportions of heteroplasmy). With higher fitness penalties *ϵ* this behaviour takes longer to evolve but still emerges.

For dynamic environments, substantially higher mutation rates, and stronger bottlenecks, evolved. Both mutation rate and bottleneck strength peaked at intermediate fitness penalties (Fig. 4A-B). Evolved bottleneck strength showed less dependence on the period of environmental change, although for very rapid environmental change bottleneck strength was marginally weaker at both low and high fitness penalties (Fig. 4B). At lower evolved mutation rates, there was a weak positive correlation between evolved mutation rate and evolved segregation strength (Fig. 4C). This correlation weakened further at high mutation rates, with evolved segregation strength plateauing around a maximum value (Fig. 4C).

The effect of the evolved nonzero mutation rates in these experiments was to maintain a low level of heteroplasmy in cells and in the population (as observed experimentally [5, 6, 49]; see Discussion), allowing more rapid adaptation when environmental change occurs. This genetic variability can be viewed as a bet-hedging mechanism, sacrificing perfect adaptation in the current state for increased flexibility in an uncertaint future [18, 19].

We also performed evolutionary simulations under fluctuating environments where we fixed mutation rate *μ* to a constant value and allowed segregation strength *σ* to evolve (Supplementary Fig. S5). We found a similar picture, with evolved segregation strength peaking at intermediate fitness penalties, and observed that stronger segregation evolved to compensate for lower mutation rates when the latter was fixed at a low value.

The evolutionary dynamics of these parameters under different environmental regimes are shown in Fig. 4D, along with example heteroplasmy distributions in a single population of cells. In all cases, as mutation rate and segregation strength increase in concert from their zero initial state, the population becomes progressively more able to adapt to the changing environments, eventually switching readily and rapidly from one environment to the other. In parallel, the population structure remains robust as before, with a high proportion of homoplasmy, even compared to the case without fluctuating environmental conditions.

## Discussion

Taken together, these results suggest that organelle heteroplasmy and bottlenecks allow an intracellular resolution to the robustness-evolvability ‘paradox’ [17] (Fig. 5). Specifically, a combination of intermediate mutation rate and segregation through a bottleneck allow organelle populations to resolve the tradeoff between robustness to mutational pressure and the ability to adapt to new environments. Low mutation rates alone allow a robust population but are insufficient to allow adaptation; high mutation rates alone allow (eventual) adaptation but lead to a less homoplasmic, less robust population. Intermediate mutation rates with segregation allow both rapid adaptation and the maintenance of robust homoplasmy in the population. We found that this combined ‘strategy’ emerges under a wide range of fluctuating environmental conditions, and that a limited level of natural heteroplasmy emerges as a consequence in evolving populations. This emergent heteroplasmy can be interpreted as a mechanism for ‘hedging bets’ against future environmental challenges [18, 19] – increasing evolvability – while its limited nature confers a degree of robustness to mutational challenge in the current environment. Heteroplasmy retains some within-cell genetic variance that can be rapidly exploited by a bottleneck in the case of an environmental shift.

**Figure 5:**
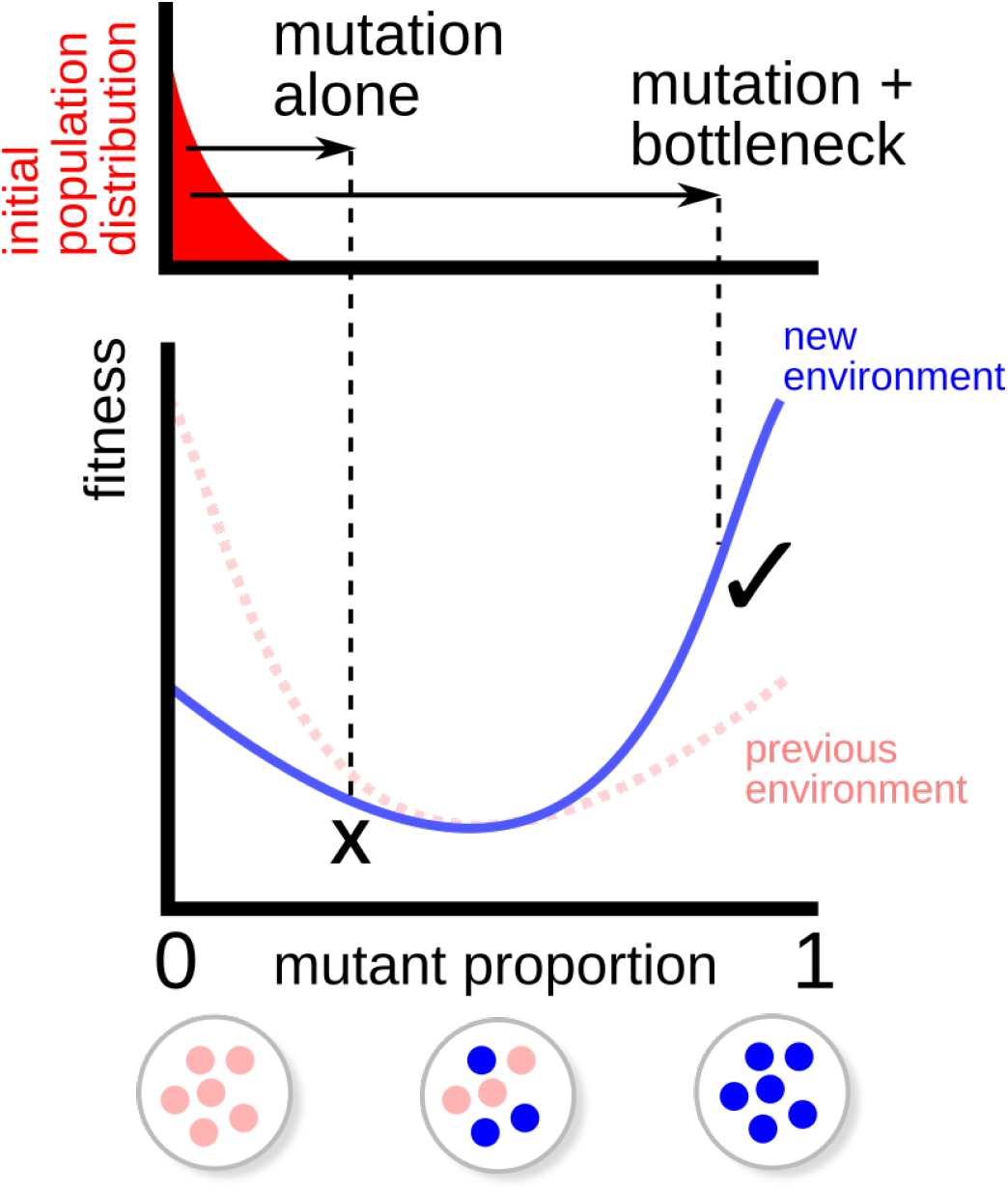
Illustration of fitness valley traversal. A population (top, red) is initially adapted to a particular environment (fitness function in dashed line), favouring a particular complement of organelle genomes (bottom). When the environment changes (fitness function in solid line), the optimal complement changes too, but the fitness penalty associated with mixed organelle populations prevents ‘hill-climbing’ adaptation to the new optimum through mutation alone (×). When combined with a genetic bottleneck, the moderate heteroplasmy arising from mutation allows some offspring to cross this fitness valley and adapt more readily to the new environment (✓).

Rather than solely being an error associated with imperfect replication and mutagen action, these results suggest that a limited amount of organelle heteroplasmy can be an evolutionarily beneficial property for systems in fluctuating conditions. Analogous results have been recently suggested for paternal leakage of organelle DNA [50], providing another source of genetic variability to facilitate environmental adaptation. In both cases, the machinery governing inheritance and maintenance of organelle genomes is itself ultimately subject to evolutonary pressures. Neglecting fundamental physical limits on the fidelity of copying and transmission [51], it may therefore be surprising if either feature was retained if it provided an exclusively negative evolutionary influence. These results suggest a possible positive influence contributing to the maintenance of these features. If biologically representative, this theory then jointly explains the presence of otherwise detrimental heteroplasmy [1] and provide a mechanism by which evolvability is itself evolvable in this system [52].

While it is often challenging to experimentally test evolutionary hypotheses, several observations are compatible with the predictions of our theory. The magnitudes of normalised heteroplasmy level variance *V*’(*h*) observed in experiments vary between genotypes and systems (with a range around 0.005 - 0.4 [27]), but are not incompatible with the *σ* ~ 0.2 - 0.4 values that emerge from our evolutionary simulation. *V*’(*h*) values around 0.2 are found, for example, in the inheritance of the disease-causing 3243 mutation in humans [53, 54, 55], for other human genotypes [56] (higher values are found for the 8993 mutation [54, 55]) and for some observations in mice [57]. Values of *V*’(*h*) around 0.1 are observed more commonly, for other human genotypes and species including insects, mice, primates, and humans [58, 59, 32, 60, 61,35]. Of course, in biology, this variance must be generated by physical mechanisms (including subsampling, turnover, and cell divisions), which may limit the ability of a system to achieve high segregation strengths.

Our model represents mutation and segregation processes through coarse-grained perturbation kernels. In reality, mutation and segregation can occur through many specific microscopic mechanisms, which previous and ongoing work has characterised in quantitative detail (reviewed in [27, 26]). We intend here to demonstrate the general principles behind this behaviour: detailed quantitative comparison between the predictions of our model and the magnitude of these effects in individual species will constitute followup work.

Regarding the qualitative predictions that our theory makes, technological advances have revealed the ubiquitous presence of mtDNA ‘microheteroplasmy’, the limited presence of a range of genetically diverse mtDNA types, in mammalian cells [5, 6]. The prediction that low-level heteroplasmy is present and inherited between generations also has experimental support in humans [49]. Rapid shifting of mtDNA types, even on the timescale of a single generation, was originally observed decades ago in cattle [42] (giving rise to the bottleneck picture). The phenomenon of ‘substoichiometric shifting’ (SSS) in plant biology provides another example [62, 63]. In SSS, mitotypes that are initially present at levels much less than one per cell in one generation can rapidly come to dominate organisms in the next generation, providing a dramatic example of fast changing mtDNA populations. Plant mtDNA undergoes frequent recombination, providing a powerful segregation effect [64] that may contribute to this rapid shifting. Our theory would suggest that the unusual strength of this capacity for segregation in plants may have evolved due to their status as sessile organisms, consequently exposed to substantial environmental fluctuations [4]. Conversely, we may expect more limited segregation to be observed in organisms less subject to environmental fluctuations.

## Acknowledgements

This project has received funding from the European Research Council (ERC) under the European Union’s Horizon 2020 research and innovation programme (Grant agreement No. 805046 (EvoConBiO) to IGJ).

## Data, code, and materials

All code is publically available at www.github.com/StochasticBiology/organelle-evolvability.

## Authors’ contributions

IGJ conceived the study, wrote the code and performed the simulations, analysed the data, drafted the manuscript, and critically revised the manuscript; ALR conceived the study, analysed the data, and critically revised the manuscript.

## Supplementary Information

**Figure S1:**
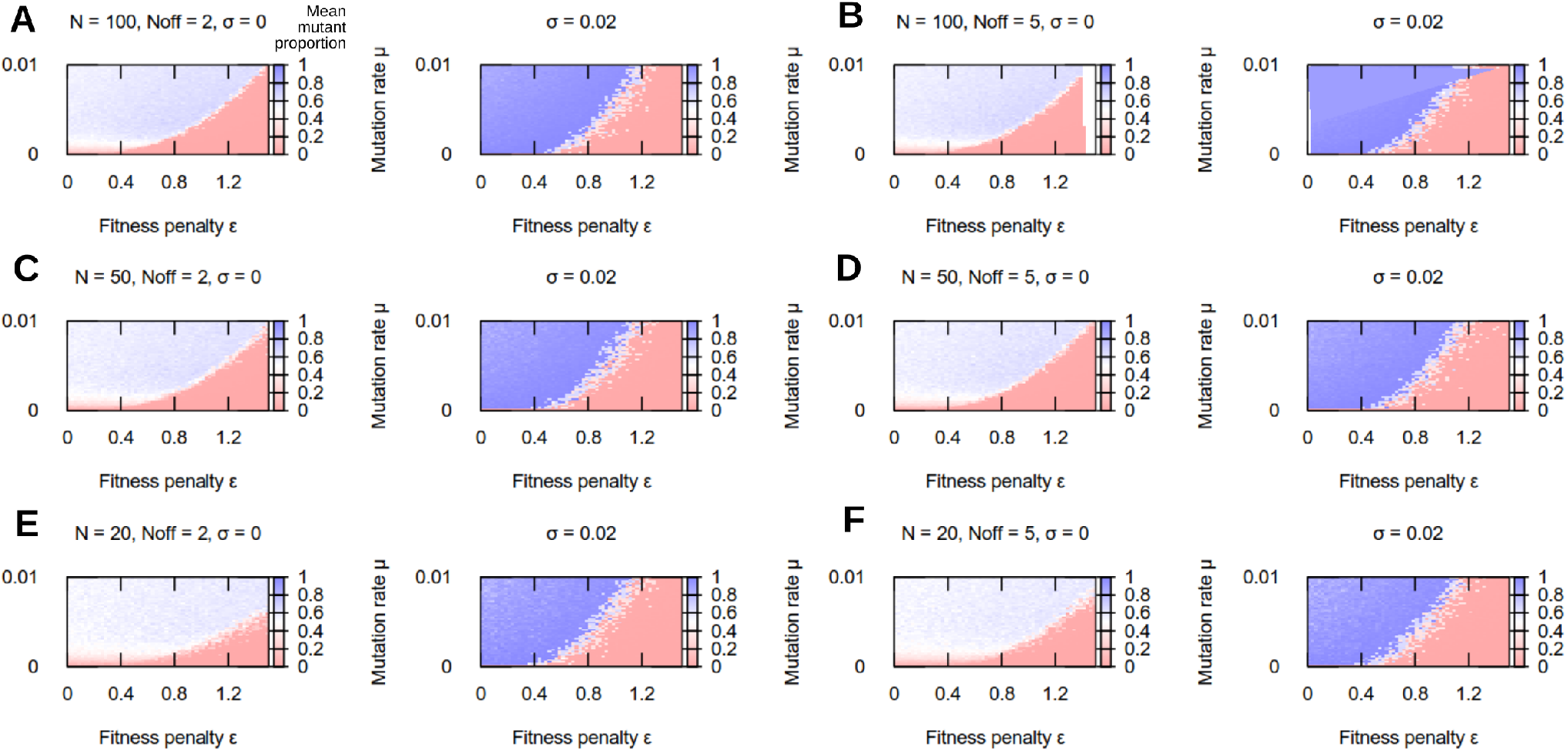
Limited influence of demographic parameters on adaptation behaviour. Mean proportion of adapted oDNAs following an environmental change, with and without segregation for different population sizes *N* and offspring numbers *N_off_*.

**Figure S2:**
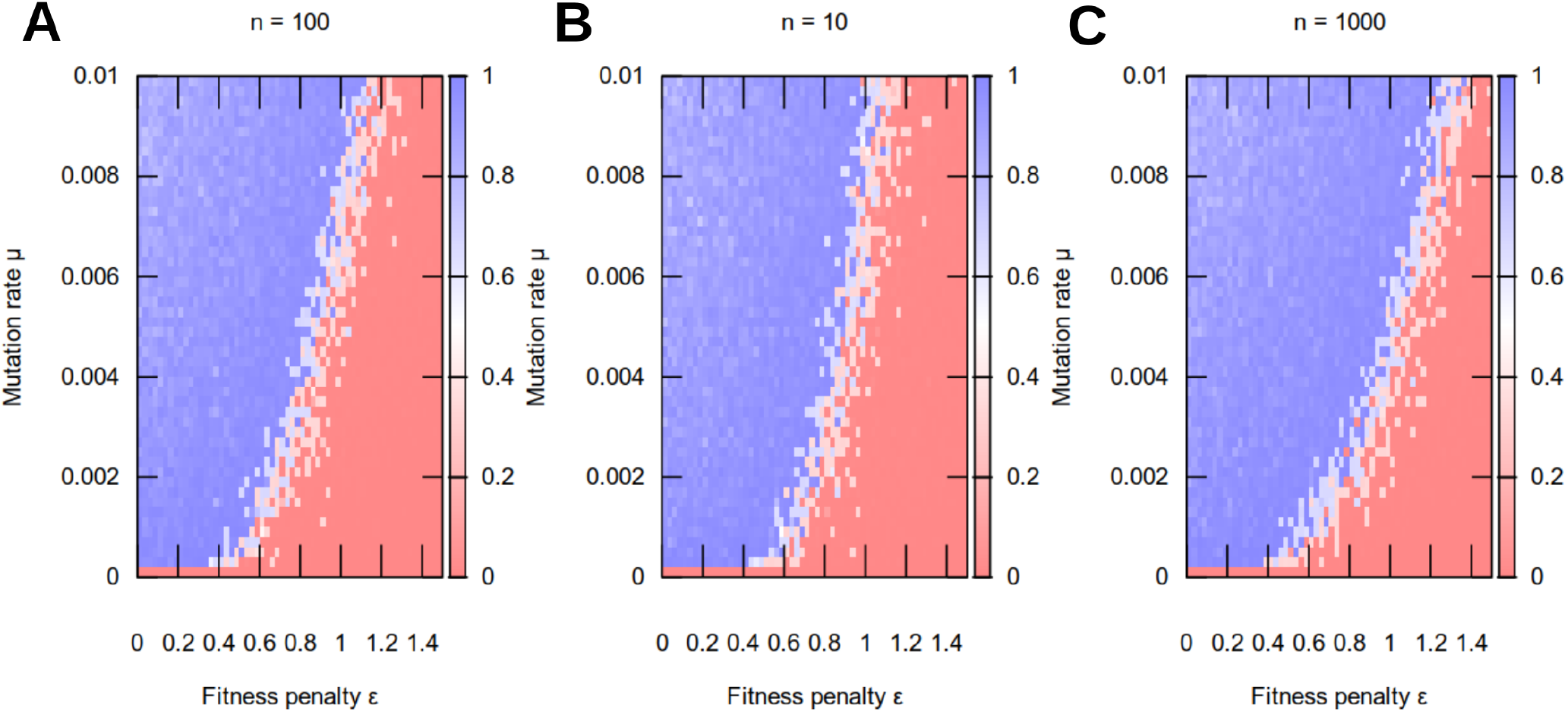
Limited influence of oDNA copy number on adaptation behaviour. Mean proportion of adapted oDNAs following an environmental change, with and without segregation for different cellular oDNA copy numbers *n*.

**Figure S3:**
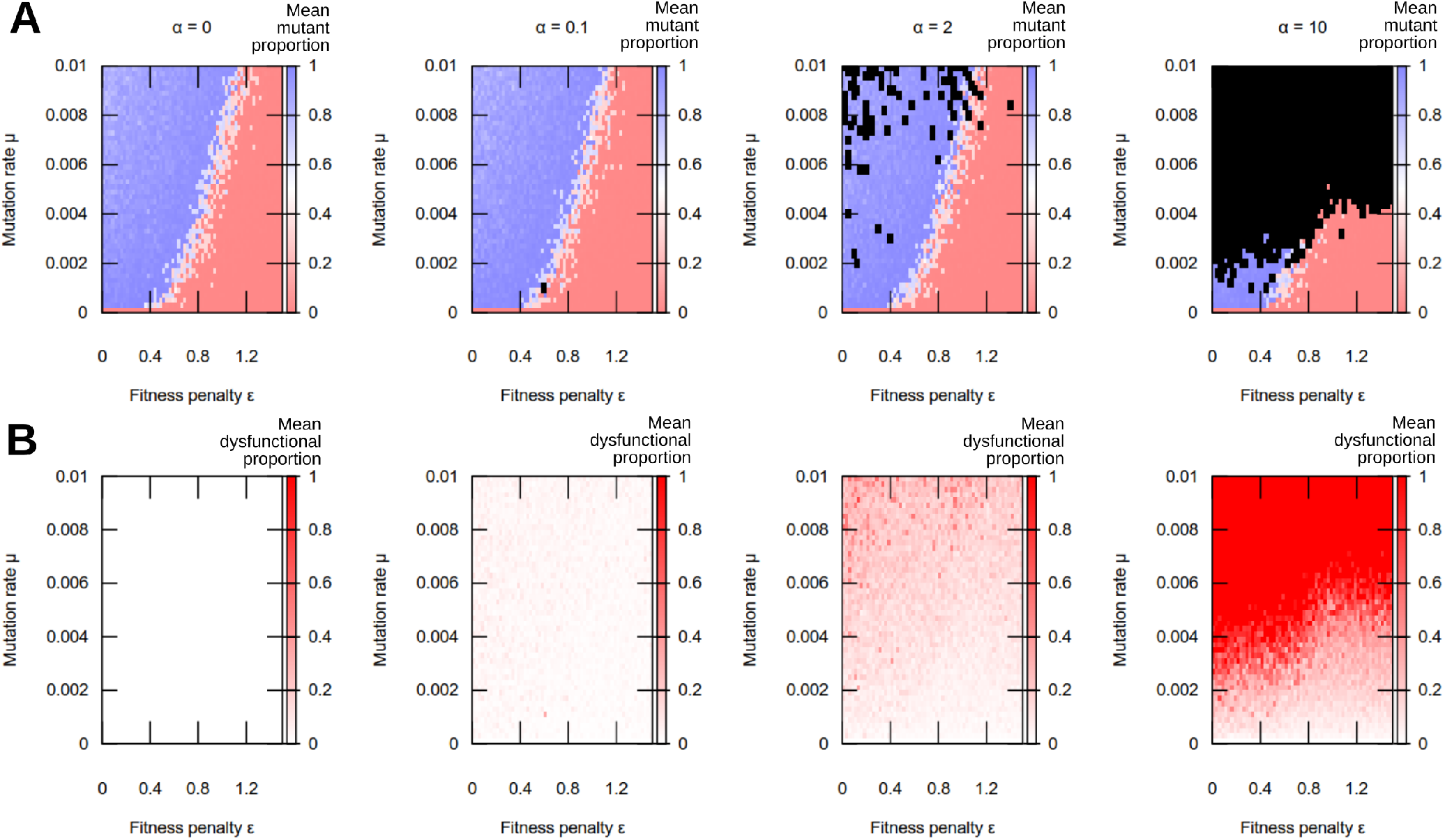
Mutations inducing dysfunctional oDNA limit feasible mutation rates but preserve adaptation behaviour. (A) Mean mutant proportion *h* and (B) mean dysfunctional proportion *h_D_* following an environmental change, with different *α*, where *μ_D_* = *αμ* is the mutation rate leading to dysfunctional oDNA. In black regions, population viability was lost (all oDNAs in a cell became dysfunctional). This viability loss appears patchily in some regions due to sampling (each pixel is the average of ten simulations; if any of these simulations lose all functional oDNA, viability loss is reported). Outside these regions, segregation and mutation facilitated adaptation as in previous cases.

### Derivation of approximate pertubation kernels for heteroplasmy statistics

We consider three random variables *W, M, D* for the copy number of wildtype, mutant, and dysfunctional oDNAs in the cell after a mutational step. If a cell currently has copy numbers *w, m, d* respectively, the effect of a mutational step is described by the following processes:

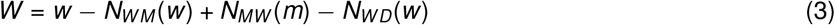

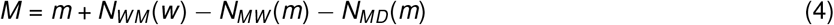

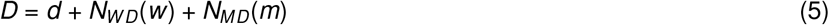

where *N_ij_*(.) are random variables describing the number of molecules of type *i* that are mutated to become molecules of type *j*. This model assumes a low mutation rate, so that double mutations are negligible. Given these definitions,

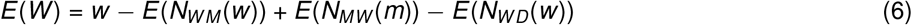

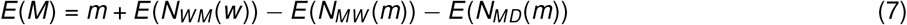

**Figure S4:**
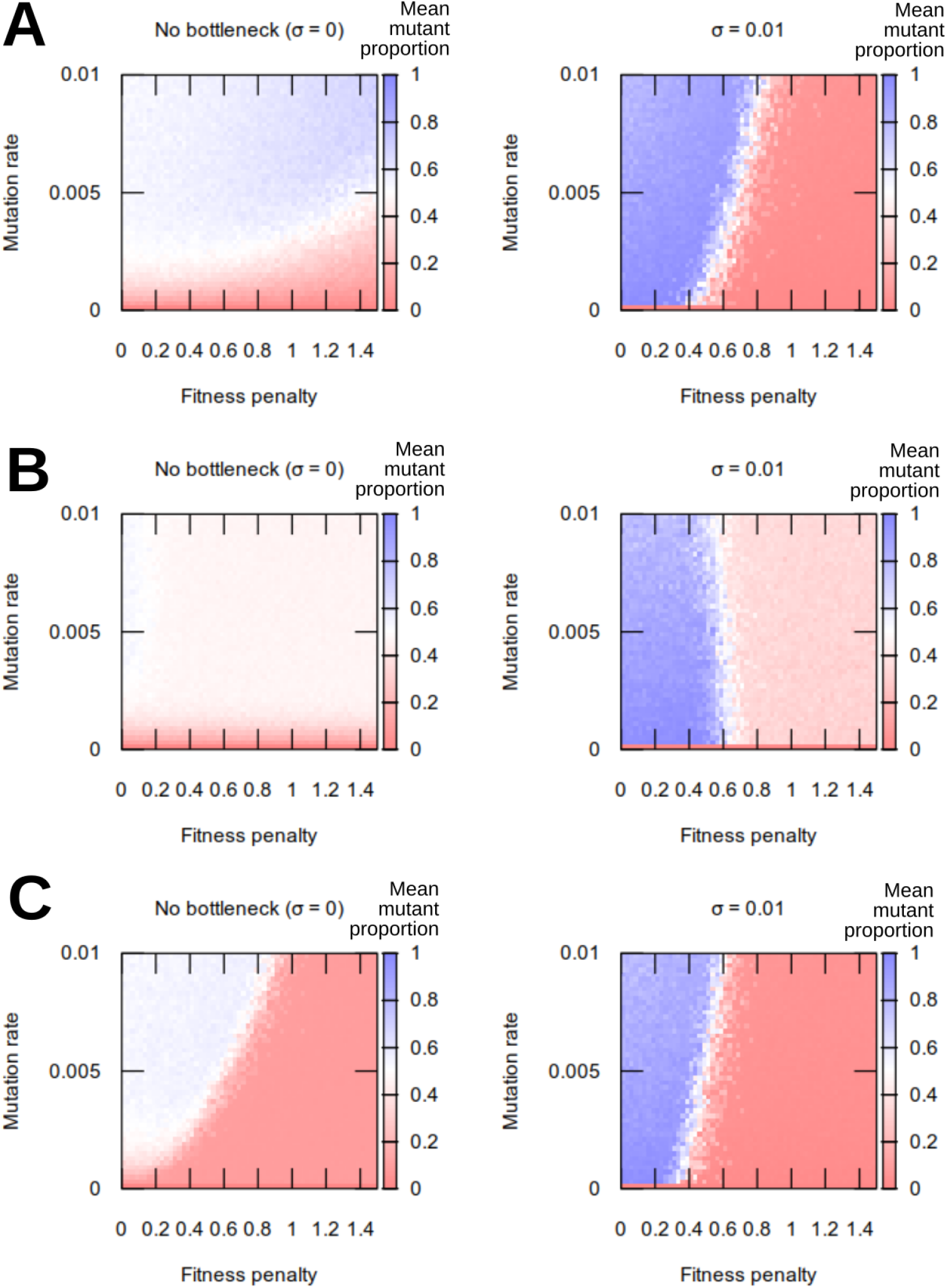
Adaptation behaviour under alternative fitness functions. Mean proportion of adapted genomes for (A) quadratic fitness penalty 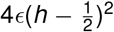; (B) large threshold (fixed penalty *ϵ* for *h* > 0.5); (C) small threshold (fixed penalty *ϵ* for *h* > 0.1). As before, adaptation is facilitated by higher segregation strength (right).

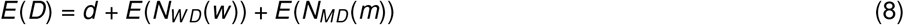

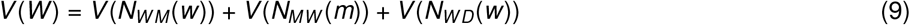

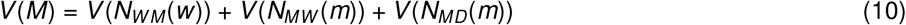

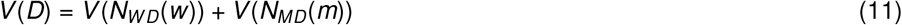

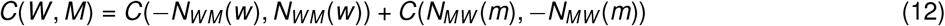

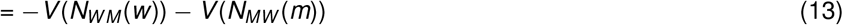

**Figure S5:**
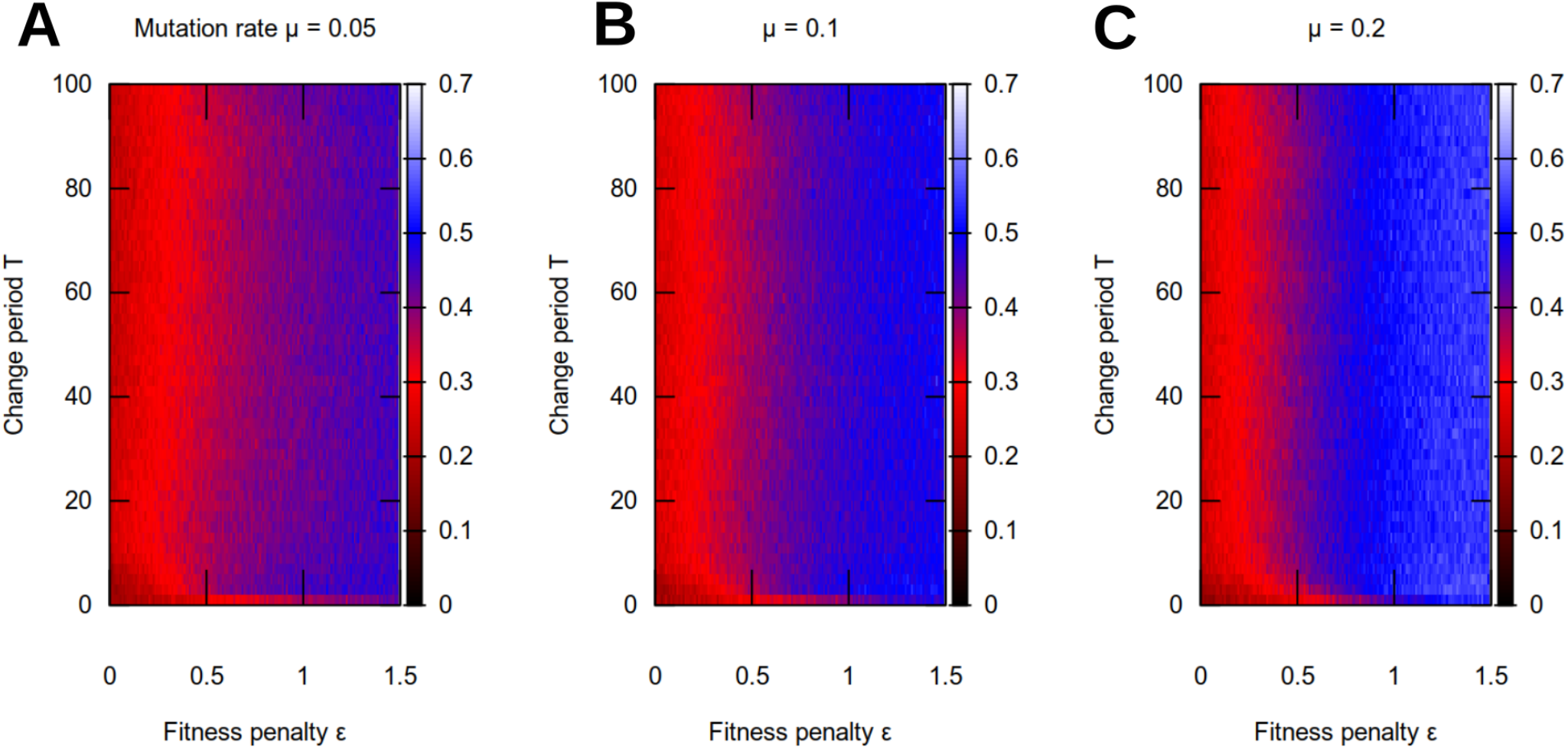
Evolution of the bottleneck under fixed mutation rates. Segregation strength *σ* emerging from evolutionary simulations (as in the main text) in fluctuating environments, for different fixed mutation rate *μ* and fitness penalty *ϵ*. At lower *μ*, stronger segregation evolves to compensate.

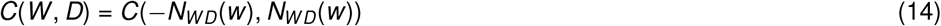

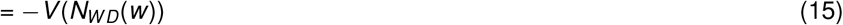

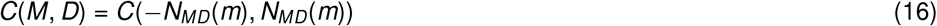

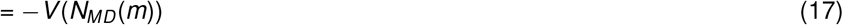

where the variance relations assume that *N_ij_* are independent. We next assume that dynamics are Poissonian:

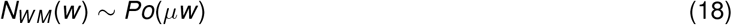

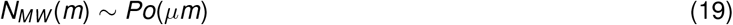

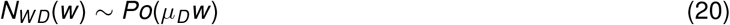

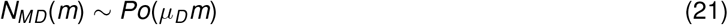

And, as *E*(*Po*(*λ*)) = *V*(*Po*(*λ*)) = *λ*,

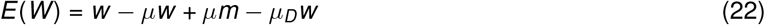

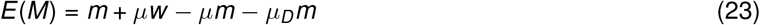

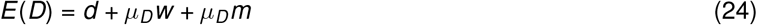

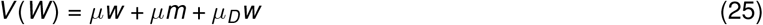

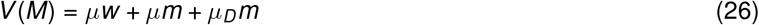

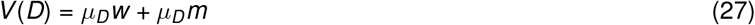

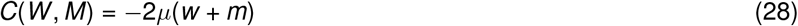

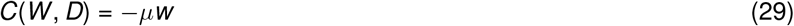

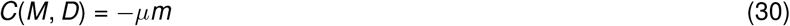

We define heteroplasmy after the mutational step by *H* = *M*/(*W* + *M*), and before the step by *h* = *m*/(*w* + *m*). First, we set *d* = 0 and *μ_D_* = 0, so that *W* + *M* = *w* + *m* = *n* remains constant, *h* = *m/n*, *w* = (1 - *h*)*n*, and *m* = *hn*. Then

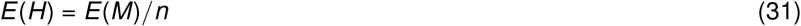

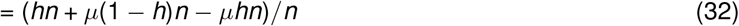

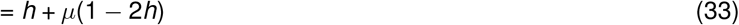

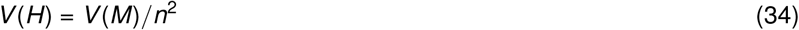

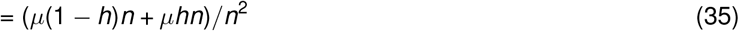

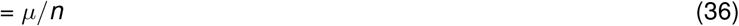

If we do not neglect d and *μ_D_*, *W* + *M* can in general depart from *w* + *m*, as functional oDNAs become dysfunctional. Now we define *N* = *W* + *M*, *H_D_* = *D/N*, *h_D_* = *d/n*, and so *w* = (1 - *h*)(1 - *h_D_*)*n, m* = *h*(1 - *h_D_*)*n, d* = *h_D_n*. We approximate moments of *H*, now being the ratio of random variables, via Taylor expansions [33]:

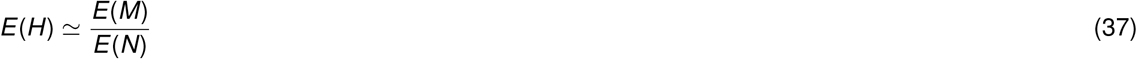

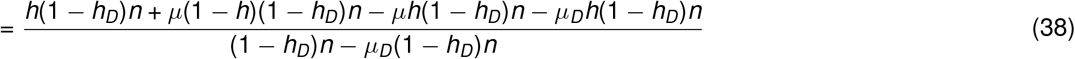

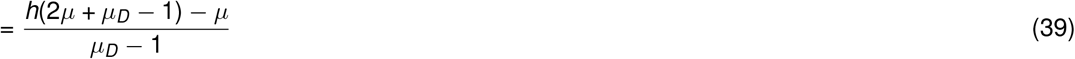

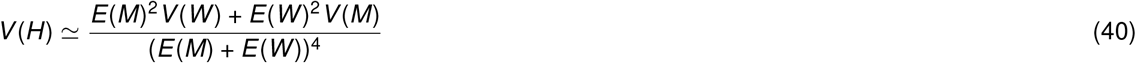

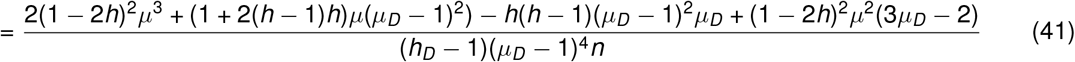

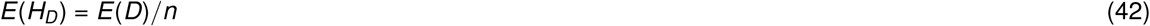

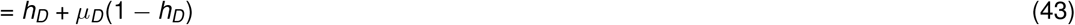

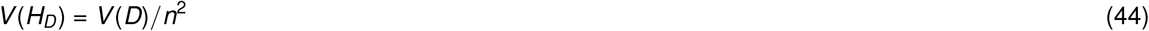

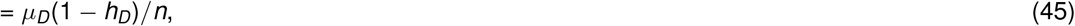

and use these moments in the normal perturbation kernels modelling the action of intergenerational mutations.

